# Fasting Rescues Locomotion in Neuromodulation-Deficient *C. elegans* via Octopamine–Gαq Signaling

**DOI:** 10.1101/2025.04.19.649495

**Authors:** Jiayi He, Guangshuo Ou, Wei Li

## Abstract

Nutrient deprivation induces adaptive behavioral and physiological changes that are critical for survival. Here, we demonstrate that fasting ameliorates locomotion defects in *Caenorhabditis elegans* mutants lacking UNC-31/CAPS, a protein essential for dense-core vesicle (DCV)-mediated neuromodulation. Through forward genetic screening, we identified a gain-of-function mutation in *egl-30*, which encodes the heterotrimeric G protein α subunit Gαq, that suppresses the locomotion defects of *unc-31* mutants under fed conditions. Transcriptomic analyses revealed that fasting induces upregulation of *egl-30* and its downstream effectors in *unc-31* mutants. Remarkably, exogenous octopamine treatment, which activates EGL-30/Gαq signaling, mimicked the fasting response and restored locomotion in an EGL-30-dependent manner. Our findings uncover a mechanism of neuromodulatory plasticity, in which metabolic stress activates a compensatory octopamine-Gαq signaling cascade to bypass impaired DCV-mediated neuromodulation, and suggest potential therapeutic strategies for CAPS-related neuropsychiatric disorders.

## Introduction

The ability of organisms to adapt behaviorally and physiologically to fluctuating nutrient availability is critical for survival. In response to nutrient deprivation, many species exhibit significant shifts in neural circuit activity, neurotransmitter signaling, and hormone release to conserve energy, promote foraging, and maintain vital motor functions (Ahn et al., 2022; Baugh and Hu, 2020; Su and Wang, 2014). Understanding the molecular basis of this adaptive response holds significant implications for deciphering how organisms modulate neural circuits and behavior in different metabolic states.

The free-living nematode *Caenorhabditis elegans* provides an ideal model to elucidate the molecular and cellular mechanisms underlying such adaptive responses, owing to its well-characterized nervous system and powerful genetic tools. Under nutrient deprivation, *C. elegans* orchestrates a fasting-induced adaptive program that includes upregulation of the bioamine octopamine, the invertebrate counterpart of norepinephrine (Rosikon et al., 2023; Suo et al., 2009; Suo et al., 2006; Tao et al., 2016). This increase is driven by the nuclear receptor DAF-12, which senses nutritional status and enhances expression of *tbh-1*, encoding the octopamine biosynthetic enzyme tyramine β-hydroxylase (Tao et al., 2016). Elevated octopamine then acts through G protein-coupled receptors (GPCR), including SER-3 and SER-6, leading to the activation of the Gαq ortholog EGL-30 (egg-laying defective family member 30), a critical regulator of neurotransmission and neurotransmitter secretion (Brundage et al., 1996; Ch’ng et al., 2008; Lackner et al., 1999; Miller et al., 1999; Suo et al., 2006; Yoshida et al., 2014). This activation, in turn, recruits its major effectors such as phospholipase Cβ (PLCβ) ortholog EGL-8 and the diacylglycerol-binding protein UNC-13/Munc13, which are essential for synaptic vesicle (SV) exocytosis and potentiates acetylcholine release at neuromuscular junctions (Augustin et al., 1999; Bastiani et al., 2003; Lackner et al., 1999; Richmond et al., 1999).

In addition to SV exocytosis, dense-core vesicle (DCV) exocytosis plays a crucial role in neuromodulation in *C. elegans* by enabling the release of neuropeptides and monoamine (Randi et al., 2023; Ripoll-Sanchez et al., 2023). This process is regulated by the single CAPS (calcium-dependent activator protein for secretion) ortholog UNC-31, which is essential for proper DCV docking and exocytosis (Ann et al., 1997; Hammarlund et al., 2008; Sieburth et al., 2007; Speese et al., 2007; Zhou et al., 2007). Loss-of-function mutations in *unc-31* impair DCV-dependent signaling, resulting in pronounced locomotion defects and an uncoordinated (Unc) phenotype (Avery et al., 1993). Notably, genetically activating the Gαs and Gαq pathway can restore locomotion in *unc-31* null mutants to near wild-type levels (Charlie et al., 2006; Zhou et al., 2007). However, it remains unclear whether physiological or environmental factors can naturally trigger compensatory mechanisms to counteract the loss of DCV-mediated neuromodulation.

Mutations in the human homolog of UNC-31, the CAPS protein family (CAPS1/CADPS1 and CAPS2/CADPS2), are implicated in a range of neurological disorders. For instance, loss of CADPS function has been linked to central nervous system primitive neuroectodermal tumors (Miller et al., 2011), while mutations in *CADPS* are strongly associated with bipolar disorder, where impaired function disrupts neuronal exocytosis, synaptic plasticity, and catecholamine uptake (Sitbon et al., 2022). Additionally, *de novo* variations in *CADPS2* have been identified in patients with autism spectrum disorder (ASD) (Bonora et al., 2014; Girirajan et al., 2013; Grabowski et al., 2017; Grove et al., 2019; Okamoto et al., 2011; Sadakata et al., 2007). Collectively, these findings suggest that defects in dense-core vesicle (DCV) exocytosis, mediated by CADPS proteins, may increase susceptibility to environmental stressors, thereby elevating the risk for neuropsychiatric conditions such as bipolar disorder and ASD. Despite these insights, effective therapeutic strategies for treating CAPS/UNC-31-associated neurological disorders remain an unmet need.

In this study, we demonstrate that fasting rescues locomotion deficits in *unc-31* mutants by activating an EGL-30/Gαq-dependent pathway. We identify a gain-of-function mutation in *egl-30* that suppresses *unc-31*-associated locomotion defects and show that fasting upregulates EGL-30/Gαq signaling components and neurotransmitter receptor genes. Exogenous octopamine treatment similarly rescues locomotion in an EGL-30/Gαq-dependent manner, linking metabolic state to neuromodulatory signaling. Our findings reveal how environmental stressors, such as fasting, can remodel neural circuits and behavior, and suggest novel therapeutic strategies for CAPS/UNC-31-associated neurological disorders.

## Results

### Fasting Rescues Locomotion Defects in *unc-31* Mutants

We found that fasting partially restores locomotion in *unc-31* mutants. In a forward genetic screen for uncoordinated (Unc) phenotypes, we isolated a *cas30000* allele of *unc-31*, which introduces a premature stop codon (Q139*) in the N-terminal coiled-coil domain of UNC-31 (Figure 1A and S1). During routine maintenance, we observed that overnight fasting unexpectedly restored motility in a subset of *unc-31(Q139*)* mutants. To systematically investigate this phenomenon, we subjected wild-type (WT) *unc-31(Q139*)* mutant animals to controlled fasting and quantified their locomotion. WT animals exhibited comparable body bending speeds under both fed and 12-hour fasting conditions (Figure 1B). In contrast, *unc-31(Q139*)* mutants displayed severe locomotion defects with complete penetrance under fed conditions, but their locomotor abilities were significantly improved after 12 hours of fasting (Figure 1C). This fasting-induced rescue was not limited to the *cas30000* allele; a similar effect was observed in another *unc-31* mutant allele, *e928* (Avery et al., 1993; Speese et al., 2007). However, fasting failed to restore locomotion in *unc-13(e51)* null mutants (Figure 1D), suggesting that the rescue specifically involves the dense-core vesicle (DCV)-mediated neuromodulation pathway. Together, these findings demonstrate that nutrient deprivation activates compensatory mechanisms to mitigate locomotion deficits caused by *unc-31* mutations.

**Figure 1.**
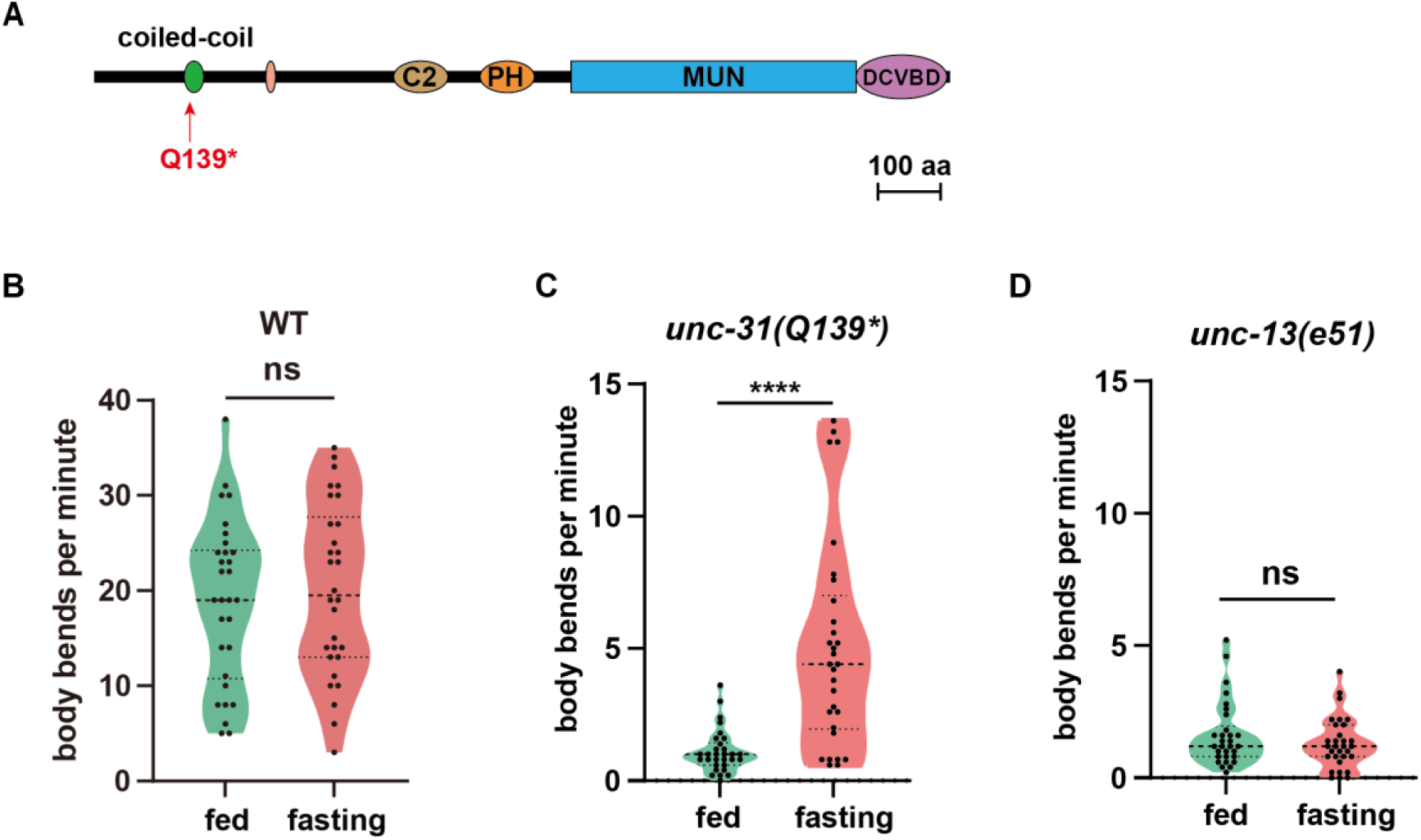
Fasting rescues locomotion of *unc-31(Q139*)* mutants. **(A)** UNC-31 protein architecture, highlighting the Q139* premature stop codon (*) in the N-terminal coiled-coil domain. **(B-D)** Violin plots showing locomotion rates of WT **(B)**, *unc-31(Q139*)* mutants **(C)** and *unc-13(e51)* mutants **(D)** under fed or 12-hour fasting conditions. Black dotted lines represent the median and quartiles. n > 30 animals. Statistical significance was calculated by unpaired t test **(B)** and unpaired Mann-Whitney test **(C and D)**.

### Genetic Screen Identifies *egl-30* Gain-of-Function Mutation as a Suppressor of *unc-31* Mutants

To elucidate the genetic mechanisms underlying the fasting-induced rescue of locomotion in *unc-31(Q139*)* mutants, we conducted a suppressor screen on fed *unc-31(Q139*)* animals (Figure 2A). Screening approximately 42,000 mutagenized haploid genomes, we isolated three suppressor alleles: *cas30038*, *cas30043*, and *cas30044*. Whole-genome sequencing and sibling subtraction analyses revealed that *cas30038* carries a dominant missense mutation (A227V) in *egl-30*, *cas30043* harbors a recessive nonsense mutation (E445*) in *Y34B4A.4*, and *cas30044* contains a deletion in *glt-1* (Figure S2A-B). Quantification of body bends demonstrated that all three suppressors significantly enhanced locomotion in *unc-31(Q139*)* mutants (Figure 2B and S2C).

**Figure 2.**
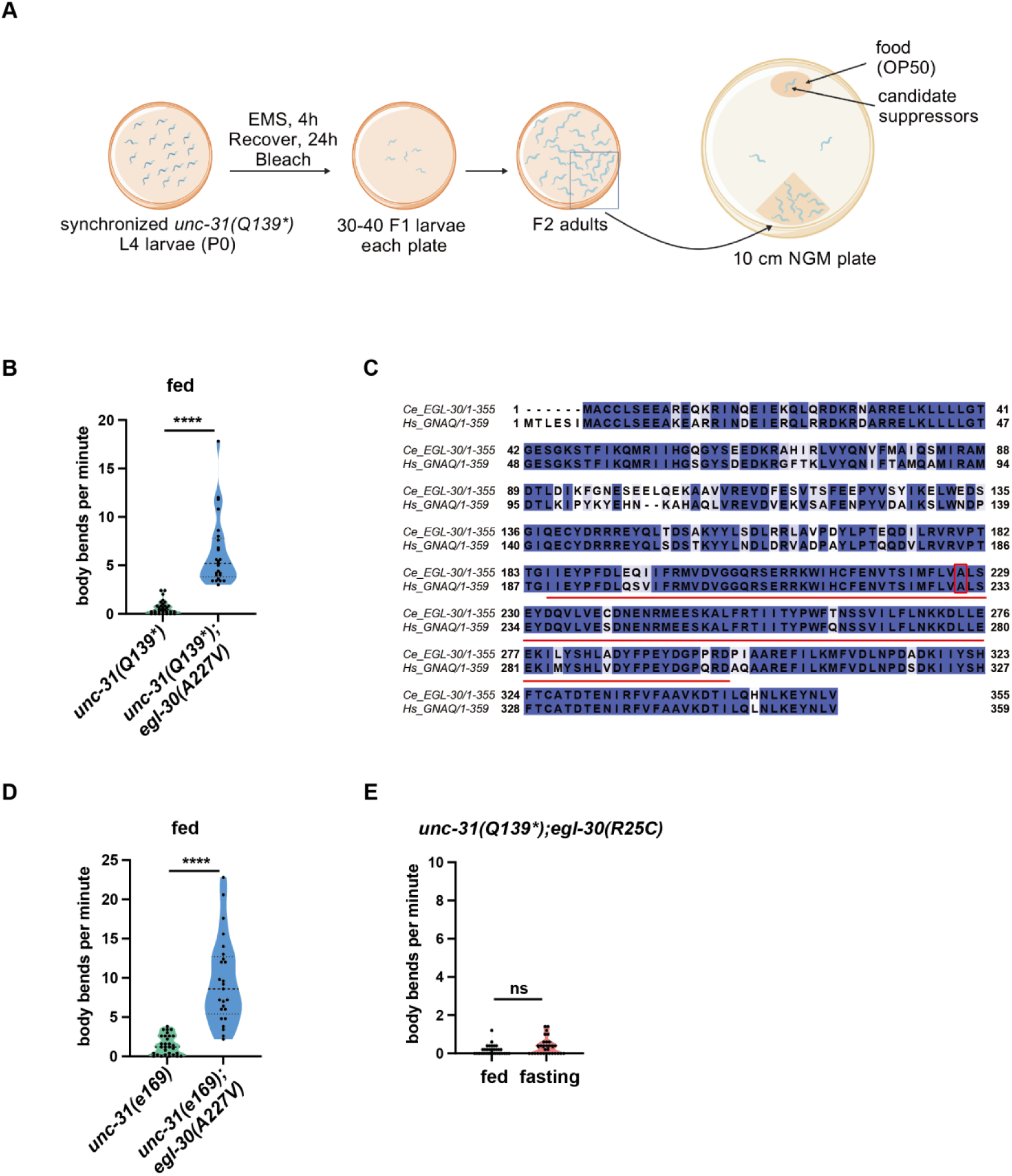
*egl-30(A227V)* rescues locomotion of *unc-31(Q139*)* mutants. **(A)** Schematic of the genetic screen for suppressors of *unc-31(Q139*)* mutant. **(B)** Violin plots showing locomotion rates of *unc-31(Q139*)* mutants and *unc-31(Q139*)*;*egl-30(A227V)* double mutants under fed conditions. Black dotted lines represent the median and quartiles. n = 25 to 30 animals. Statistical significance was calculated by unpaired Mann-Whitney test. ****P < 0.0001. **(C)** Sequence alignment of human GNAQ and *C. elegans* EGL-30 proteins. The red box highlights the conserved A227 in EGL-30 and A231 in GNAQ and the red lines highlight GTPase activity domain. **(D)** Violin plots showing locomotion rates of *unc-31(e169)* mutants and *unc-31(e169)*;*egl-30(A227V)* double mutants under fed conditions. Black dotted lines represent the median and quartiles. n = 25 to 30 animals. Statistical significance was calculated by unpaired Mann-Whitney test. ****P < 0.0001. **(E)** Violin plots showing locomotion rates of *unc-31(Q139*)*;*egl-30(R25C)* double mutants under fed or 12-hour fasting conditions. Black dotted lines represent the median and quartiles. n = 30 to 32 animals. Statistical significance was calculated by unpaired Mann-Whitney test. ns, not significant.

Among these suppressors, we focused on *egl-30*, which encodes the Gαq subunit orthologous to human GNAQ. The A227 residue lies within the conserved GTPase domain of EGL-30/GNAQ (Kamato et al., 2017; Oldham and Hamm, 2008) (Figure 2C), suggesting that the dominant A227V mutation may result in constitutive activation of EGL-30. This aligns with previous findings that hyperactivation of the Gαq pathway enhances locomotion in *unc-31* null mutants (Charlie et al., 2006). Supporting its role as a suppressor, the *egl-30(A227V)* mutation also partially rescued locomotion defects in the *unc-31(e169)* allele (Figure 2D). To further validate the functional significance of EGL-30/Gαq activity, we examined the effect of a loss-of-function allele, *egl-30(R25C)*, previously identified in our laboratory (Ke et al., 2025). In contrast to the gain-of-function *egl-30(A227V)* mutation, *egl-30(R25C)* failed to rescue locomotion in *unc-31(Q139*)* mutants. Instead, *unc-31(Q139*);egl-30(R25C)* double mutants exhibited exacerbated paralysis (Figure 2E), underscoring the necessity of upregulated EGL-30/Gαq activity for rescuing *unc-31* locomotion defects. These genetic data collectively demonstrate that enhanced EGL-30/Gαq signaling is critical to compensate for the locomotion deficits caused by *unc-31* mutations.

### Transcriptomic Profiling Reveals Upregulation of Genes Associated with the EGL-30/Gαq Pathway

To further elucidate the molecular mechanisms underlying fasting-induced locomotion recovery in *C. elegans*, we performed RNA sequencing on synchronized day 1 adult WT and *unc-31(Q139*)* mutants under fed conditions, or after 6 and 12 hours of fasting (Figure 3A). Principal component analysis (PCA) revealed distinct gene expression profiles between fed and fasted animals, with greater divergence observed after 12 hours of fasting (Figure S3). This temporal divergence suggests that prolonged fasting induces more pronounced transcriptional changes.

**Figure 3.**
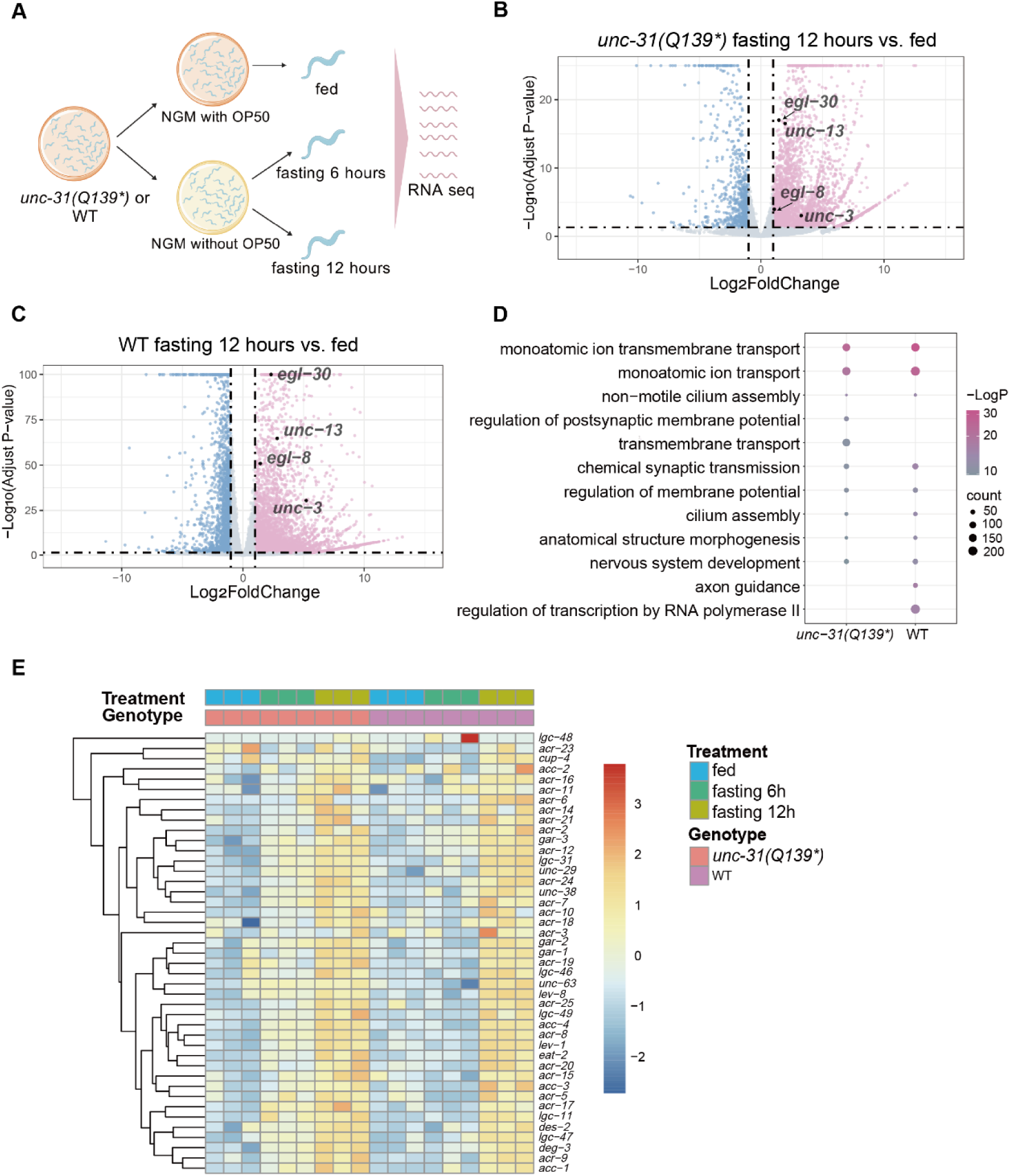
Transcriptomic profiling reveals that 12-hour fasting induced the upregulation of genes associated with the EGL-30/Gαq pathway. **(A)** Experimental workflow of RNA-seq in fed and fasted WT and *unc-31(Q139*)* mutants. **(B, C)** Volcano plots of deferentially expressed genes in *unc-31(Q139*)* mutants **(B)** and WT animals **(C)** after 12-hour fasting. Pink and blue dots represent genes that are significantly upregulated and downregulated, respectively (|fold change| > 2, adjusted P-value < 0.05). For clarity, points with -Log_10_(Adjusted P-value) > 25 **(B)** or > 100 **(C)** were capped at 25 and 100, respectively. **(D)** Dot plot of Gene Ontology (GO) enrichment analysis for biological processes in genes upregulated after 12-hour fasting in *unc-31(Q139*)* and WT strains. Dot colors indicate P-values and dot size corresponds to the number of genes enriched in each biological process. **(E)** Heatmap of expression levels of genes encoding acetylcholine receptors. The list of receptor genes was obtained from WormAtlas (https://www.wormatlas.org/NTRmainframe.htm).

To identify key pathways activated during fasting, we analyzed genes upregulated after fasting (fold change > 2, adjusted *P*-value < 0.05) (Figure 3B-E, S4, S5 and S6). Under fed conditions, *unc-31* mRNA expression levels in *unc-31(Q139*)* mutants were comparable to WT, indicating that the nonsense mutation does not affect *unc-31* mRNA levels and likely results in a truncated protein (Figure S5A). Interestingly, fasting enhanced *unc-31* expression in both *unc-31(Q139*)* mutants and WT animals (Figure S5A), suggesting a potential feedback mechanism in response to nutrient deprivation. Consistent with the genetic evidence that activation of the EGL-30/Gαq pathway by the *egl-30(A227V)* mutation rescues *unc-31(Q139*)* locomotion defects, 12-hour fasting upregulated *egl-30*, its downstream effectors *egl-8* (encoding PLCβ), and *unc-13* in both WT and *unc-31(Q139*)* mutants (Figure 3B-C and S5B-D). These findings suggest that fasting restores locomotion in *unc-31(Q139*)* mutants by upregulating genes associated with the EGL-30/Gαq pathway, thereby enhancing synaptic signaling and neuronal function.

Gene Ontology (GO) analysis of upregulated genes highlighted significant enrichment of pathways involved in synaptic function and neurotransmitter signaling (Figure 3D and S4C). This included genes encoding receptors for acetylcholine, glutamate, and GABA, as well as the transcription factor *unc-3*, which regulates cholinergic motor neuron identity and the expression of acetylcholine receptor subunit genes (Kratsios et al., 2011) (Figure 3E, S5E and S6). Notably, the 12-hour fasting group exhibited more pronounced upregulation than the 6-hour fasting group, indicating a time-dependent transcriptional response to nutrient deprivation. These results suggest that fasting activates a coordinated transcriptional program that enhances both presynaptic neurotransmitter release and postsynaptic sensitivity, thereby facilitating locomotion recovery in *unc-31(Q139*)* mutants, highlighting the interplay between metabolic state, synaptic function, and neuronal signaling.

### Octopamine Mimics Fasting to Rescue Locomotion via EGL-30/Gαq Signaling

To test whether octopamine mediates fasting-induced locomotion recovery, we treated fed *unc-31(Q139*)* mutants with octopamine. Behavioral assays showed that octopamine partially rescued locomotor defects in these mutants under fed conditions (Figure 4A). This rescue was EGL-30-dependent, as *unc-31(Q139*);egl-30(R25C)* double mutants exhibited no improvement with octopamine treatment (Figure 4B). These findings indicate that octopamine activates the EGL-30/Gαq pathway to restore locomotion, mirroring the fasting response. Together, our results establish a molecular link between nutrient deprivation and locomotion recovery through octopamine-mediated EGL-30/Gαq signaling.

**Figure 4.**
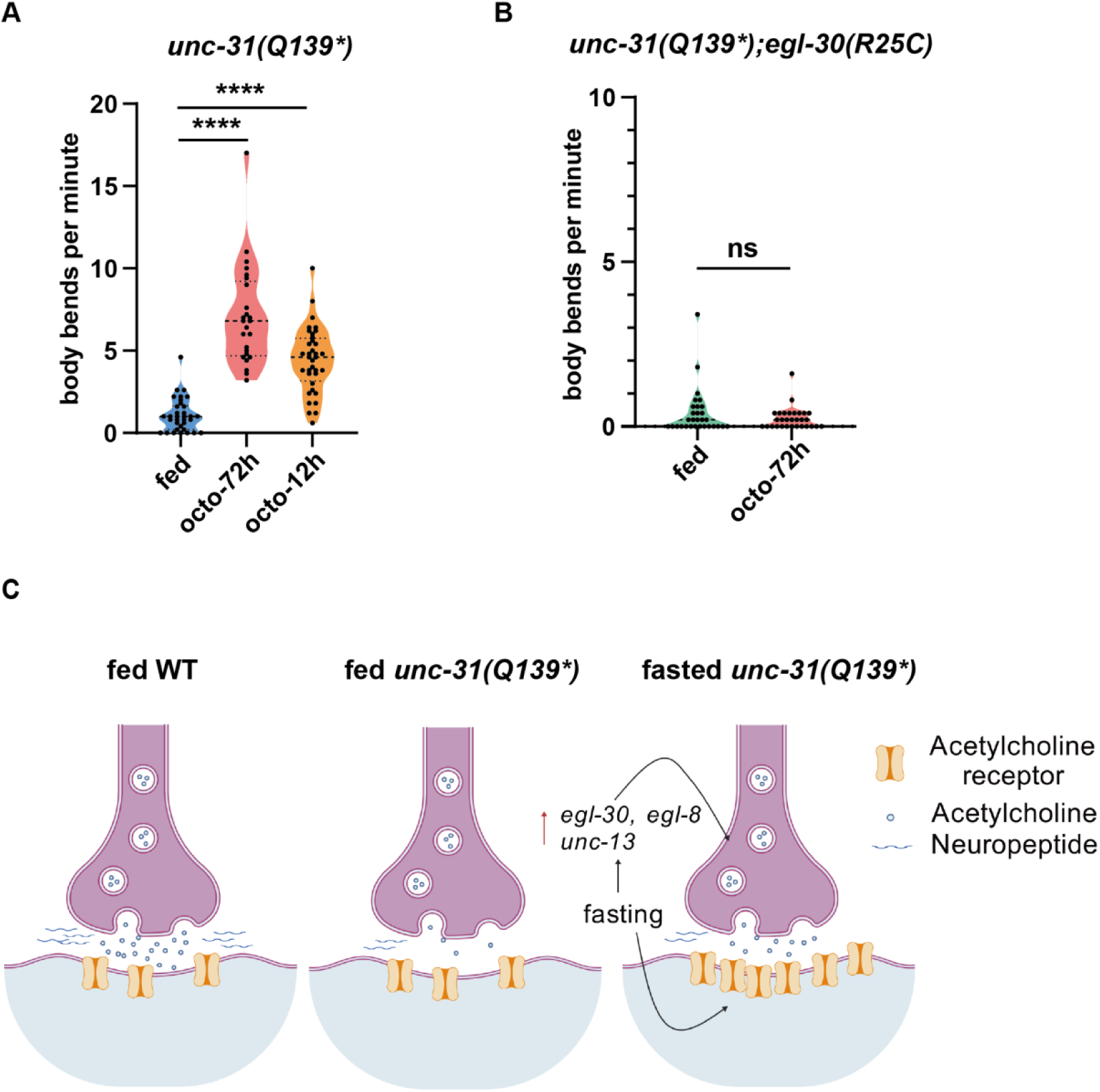
Octopamine rescues *unc-31* locomotion via EGL-30 signaling and a proposed model. **(A-B)** Violin plots showing locomotion rates of day 1 adult *unc-31(Q139*)* mutants **(A)** and *unc-31(Q139*);egl-30(R25C)* double mutants **(B)** under normal feeding conditions (fed) or 10mg/ml octopamine treatment. The octo-72h group was exposed to octopamine from L1 to day 1 adulthood (72 hours total), while the octo-12h group received treatment starting at day 1 of adulthood for 12 hours. Black dotted lines represent the median and quartiles. n =25 to 35 animals. Statistical significance was calculated by unpaired Mann-Whitney test. ****P < 0.0001; ns, not significant. **(C)** A proposed model for fasting-induced rescue of *unc-31* locomotion defects. Loss of UNC-31/CAPS impairs dense-core vesicle (DCV)-mediated acetylcholine (ACh) release. Fasting triggers two compensatory mechanisms: (1) Presynaptic enhancement: Upregulation of genes involved in ACh synthesis (e.g., *unc-17*, *cha-1*) and secretion (e.g., *egl-30/Gαq*, *unc-13*); and (2) Postsynaptic priming: Increased expression of ACh receptor subunits (e.g., *unc-29*, *unc-38*). This dual regulation restores synaptic transmission and neuromuscular function, rescuing locomotion in *unc-31* mutants.

## Discussion

Our study reveals that fasting—a common environmental stressor—activates a compensatory molecular mechanism in *C. elegans* that restores locomotion in *unc-31* mutants lacking dense-core vesicle (DCV)-mediated neuromodulation (Figure 4C). Central to this adaptation is the EGL-30/Gαq signaling pathway, as evidenced by the rescue of locomotion through a gain-of-function *egl-30* mutation and fasting-induced upregulation of *egl-30* and its downstream effectors. These findings demonstrate how organisms leverage environmental cues to mitigate genetic defects, underscoring the dynamic interplay between metabolism and neural function.

Transcriptomic profiling revealed that fasting triggers a coordinated upregulation of synaptic signaling components, including *egl-30*, its effector *egl-8/PLCβ*, and neurotransmitter receptors. This transcriptional remodeling suggests a state of heightened neural plasticity, where enhanced presynaptic secretion and postsynaptic sensitivity compensate for UNC-31 deficiency. Such adaptability may reflect an evolutionarily conserved strategy to sustain critical behaviors under stress. For instance, nutrient deprivation in *Drosophila* similarly increases locomotion to promote food-seeking (Koon et al., 2011; Yang et al., 2015; Yu et al., 2016). Whether analogous mechanisms exist in mammals, particularly in contexts of metabolic or neurological dysfunction, warrants further investigation.

Octopamine, the invertebrate counterpart of norepinephrine, emerged as a key mediator of this adaptive response. Exogenous octopamine restored locomotion in *unc-31* mutants via EGL-30/Gαq signaling, mirroring the fasting response. This highlights neuromodulators as critical conduits linking metabolic state to neural circuit plasticity. Given the functional parallels between octopamine and norepinephrine, our findings suggest therapeutic potential for targeting norepinephrine signaling in human CAPS-associated disorders. Mutations in the CAPS family (*CAPS1/CADPS1* and *CAPS2/CADPS2*) are linked to neuropsychiatric conditions such as bipolar disorder, autism spectrum disorder (ASD), and neuroectodermal tumors, where DCV exocytosis defects may heighten sensitivity to environmental stressors (Bonora et al., 2014; Girirajan et al., 2013; Miller et al., 2011; Sitbon et al., 2022). Dysregulated catecholamine signaling, a hallmark of these disorders, could be modulated by augmenting norepinephrine pathways or downstream effectors like Gαq.

In summary, our work uncovers a fasting-induced mechanism that bypasses DCV dysfunction via octopamine-EGL-30/Gαq signaling, offering insights into neural adaptability and therapeutic strategies for CAPS-related neurological disorders. Future studies should explore whether metabolic interventions or neuromodulator-targeted therapies can similarly restore synaptic function in mammals, particularly in conditions marked by synaptic vesicle trafficking deficits.

## MATERIALS AND METHODS

### Worm strains and culture

Strains were maintained on Nematode Growth Medium (NGM) plates seeded with *E. coli* OP50 at 20℃, following the standard protocols (Brenner, 1974). The wild-type strain was Bristol N2. All the *C. elegans* strains used in this study are summarized in Table S1.

### Locomotion assay

The locomotion rate was quantified by counting the body bends of Day 1 adult animals. Synchronized L1 larvae were placed on NGM plates with OP50 and grown at 20℃ until they matured into young adults. For the fasting treatment, animals were washed three times to remove OP50, then placed on NGM plates without OP50 for 12 hours. Locomotion was recorded for 5 minutes. A body bend was defined as the head or tail tip passing through maximum or minimum amplitude. The average number of body bends per minute was calculated as the total number of body bends in 5 minutes divided by 5, and served as a quantitative indicator. The graphing and statistical analysis were performed using GraphPad Prism 9.

### EMS mutagenesis and suppressor screen

The *unc-31(Q139*, cas30000)* mutant animals (P0 generation) were synchronized at the late L4 larval stage, collected in 4 mL of M9 buffer, and incubated with 50 mM ethyl methanesulfonate (EMS) for 4 hours at room temperature while being continuously rotated. After the incubation, the animals were washed three times with M9 buffer and cultured under standard laboratory conditions. Twenty-four hours later, the adult P0 animals were subjected to the bleaching process. The L1 larval F1 generation progeny were then distributed and raised on 6-cm NGM plates, with each plate containing 30-40 larvae.

For screening, 10-cm NGM plates were prepared at least one day in advance of examining the F2 generation animals. These screening plates had only a small portion of the edges seeded with *E. coli* OP50, while the rest of the area remained unseeded. Adult F2 animals were transferred to the opposite side of the OP50 lawn on these screening plates. Basically worms that were able to move to the OP50 lawn (the opposite margin of the plates) within 2 hours were very likely to be suppressors.

After two rounds of screening, approximately 42,000 haploid genomes were examined, and suppressors from different F1 plates were considered independent. After phenotype confirmation over two generations, suppressors were sent to whole genome sequencing followed by bioinformation analyses and sibling subtraction analyses to identify the genomic mutations responsible for the rescue.

### Sibling Subtraction Analyses

Sibling subtraction was performed as described before (Joseph et al., 2018) with some modifications. *unc-31(Q139*)* hermaphrodites were crossed with *him-5* males. The male progeny resulting from this cross were then selected and mated with the suppressor hermaphrodites. The F1 progeny were isolated into individual plates and subjected to genotyping to confirm the presence of the *unc-31(Q139*)* allele. The *unc-31(Q139*)* heterozygotes were then maintained, and a total number of about 40 F2 self-progeny were cloned out onto individual plates to obtain homozygous suppressor and non-suppressor strains. These strains were then mixed separately and sent for whole genome sequencing. The requirement for variant calls from the suppressed DNA pools was 90%, while the requirement for nonsuppressed variant reads was 10%.

### RNA sequencing

The “fed” and “fasted” worms were picked into 0.2 ml TRIzol and sent to mRNA sequencing. Three batches of worms were collected for treatment. The resulting raw reads of RNA-seq were assessed for quality, adaptor content, and duplication rates with FastQC. The raw sequencing reads were trimmed using Trim_galore (version 0.6.10) to remove the low-quality bases and adaptor sequences. Paired-end reads were aligned to *C. elegans* reference genome (WBcel235) using HISAT2 (version 2.2.1). The numbers of reads aligned to genes were quantified by featureCounts (version 2.0.6) in a strand specific manner. Gene names were annotated, and differentially expressed genes were identified using the OmicVerse package with DESeq method in the Python programming language. Differentially expressed genes were defined with the following criteria: upregulated genes (false discovery rate (FDR) less than 0.05, log2-transformed fold change greater than 1); downregulated genes (FDR less than 0.05, log2-transformed fold change less than -1). DAVID website (https://davidbioinformatics.nih.gov/) was used to analyze gene enrichment terms in upregulated or down-regulated genes. The GO dot plots were plotted by https://www.bioinformatics.com.cn (last accessed on 10 Dec 2024), an online platform for data analysis and visualization. The volcano plots, PCA plots and heatmaps were generated using the R packages of ggplot2, DESeq2 and pheatmap, respectively. Experimental workflow of RNA-seq (Figure 3A) and other mechanism diagrams (Figure 2A and 4C) were plotted with BioGDP.com (Jiang et al., 2025).

### Octopamine Treatment Assay

To administer octopamine hydrochloride ( MCE, HY-B0528A), the drug was added to the NGM medium at 10 mg/ml prior to pouring the 3.5-cm plates, which were subsequently seeded with OP50 culture in the usual manner.

For the 72-hour octopamine treatment, L1 larvae were distributed and raised on NGM plates containing octopamine hydrochloride. After 72 hours, the Day 1 adults were counted for body bends. For the 12-hour octopamine treatment, synchronized Day 1 young adults, raised under standard conditions, were distributed on 3.5-cm NGM plates with octopamine hydrochloride. After 12 hours, the treated worms were counted for body bends.

## Acknowledgments

This work was supported by the National Key R&D Program of China Grants 2022YFA1302700, 2024YFA1307301, 2019YFA0508401; National Natural Science Foundation of China Grants 32270773, 32470730, 32070706, 92254306, 31991190 and 32270721; Beijing Natural Science Foundation Grants QY23137; and Tsinghua University Initiative Scientific Research Program (Student Academic Research Advancement Program).

---

**Figure S1.**
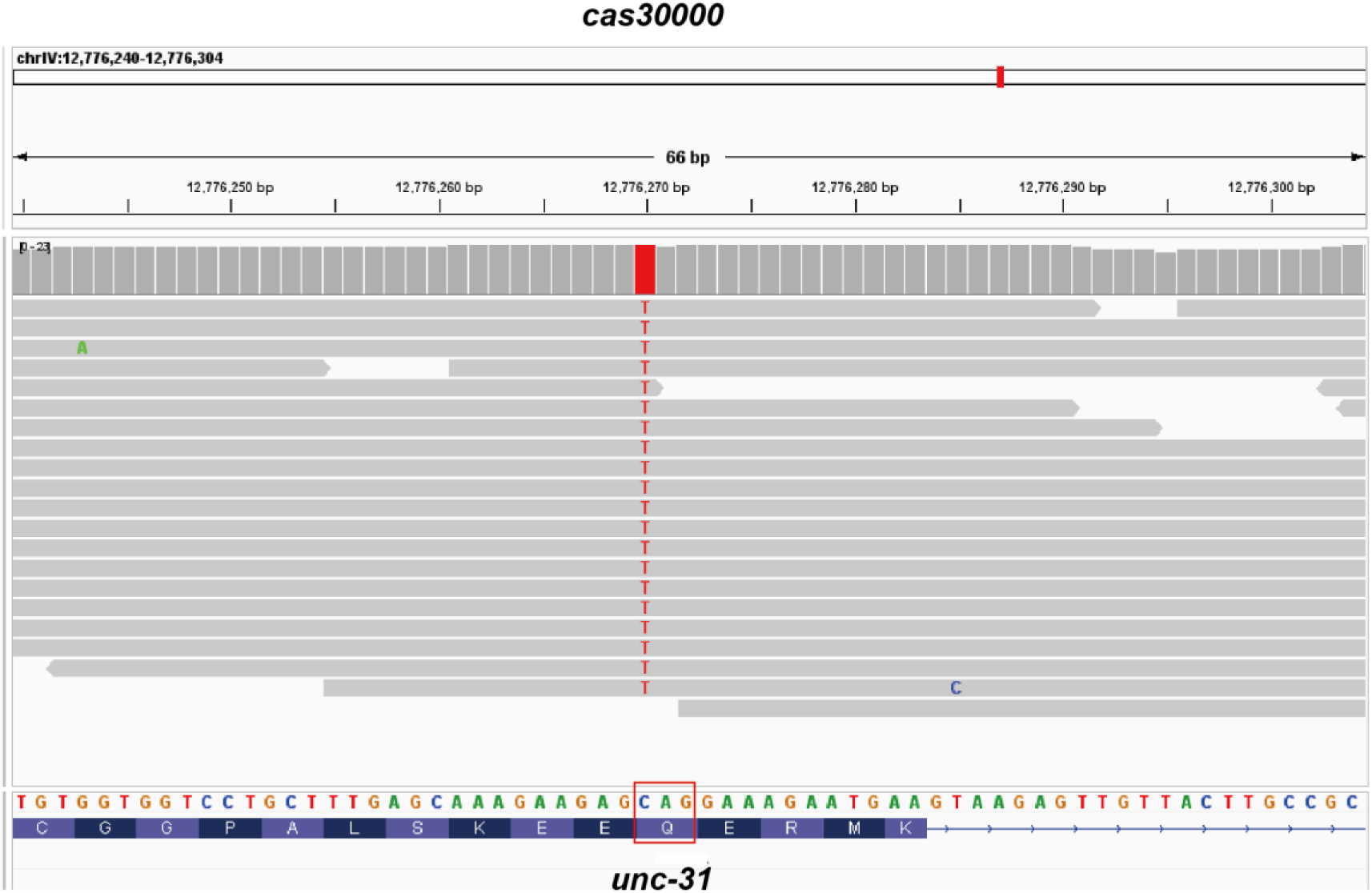
Genome sequencing of *cas30000* allele reveals a nonsense mutation in *unc-31*. IGV visualization of genome sequencing of homozygous *cas30000 at unc-31* locus. The lower panel shows the reference genome sequence and the translated amino acid sequence. The red box highlights the nonsense mutation (CAG to TAG) in *unc-31*.

**Figure S2.**
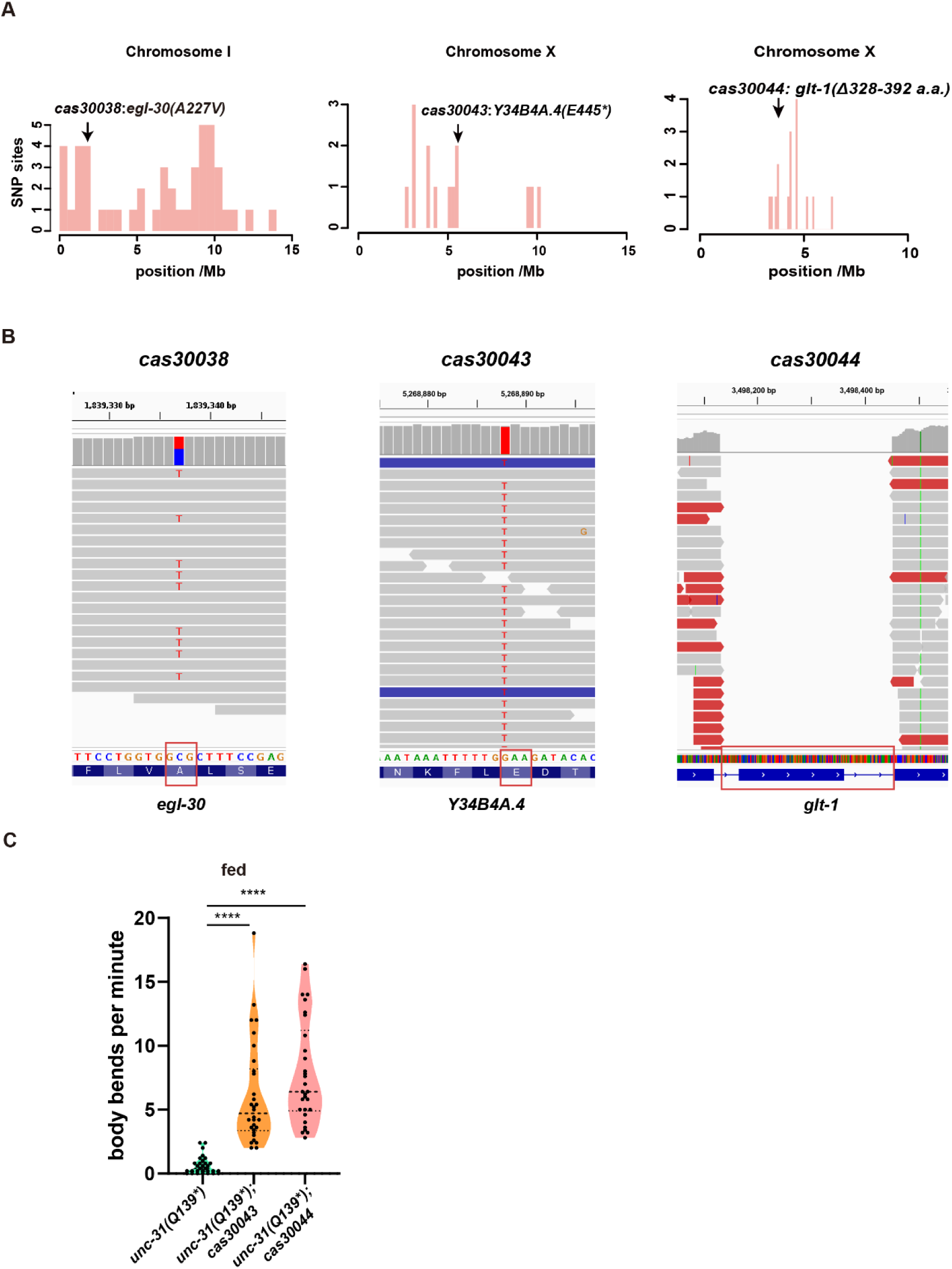
Identification of three suppressor mutations of *unc-31(Q139*)*. **(A)** EMS density mapping of *cas30038*, *cas30043* and *cas30044* suppressors of *unc-31(Q139*)*. X-axis: Chromosomal position (Mb). Y-axis: number of EMS signature SNPs. **(B)** IGV visualization of whole genome sequencing of *cas30038*, *cas30043*, and *cas30044 at egl-30, Y34B4A.4* and *glt-1* loci, respectively. The lower panel shows the reference genome sequence and the translated amino acid sequence, or the gene structure. The red box highlights the missense mutation (GCG to GTG, A227V) in *egl-30*, the nonsense mutation (GAA to TAA, E445*) in *Y34B4A.4*, the deletion of the 6^th^ exon (Δ328-392 amino acids) in *glt-1*. **(C)** Violin plots showing locomotion rates of *unc-31(Q139*)* mutants, *unc-31(Q139*)*;*cas30043* and *unc-31(Q139*)*;*cas30044* double mutants under fed conditions. Black dotted lines represent the median and quartiles. n > 25 animals. Statistical significance was calculated by unpaired Mann-Whitney test. ****P < 0.0001.

**Figure S3.**
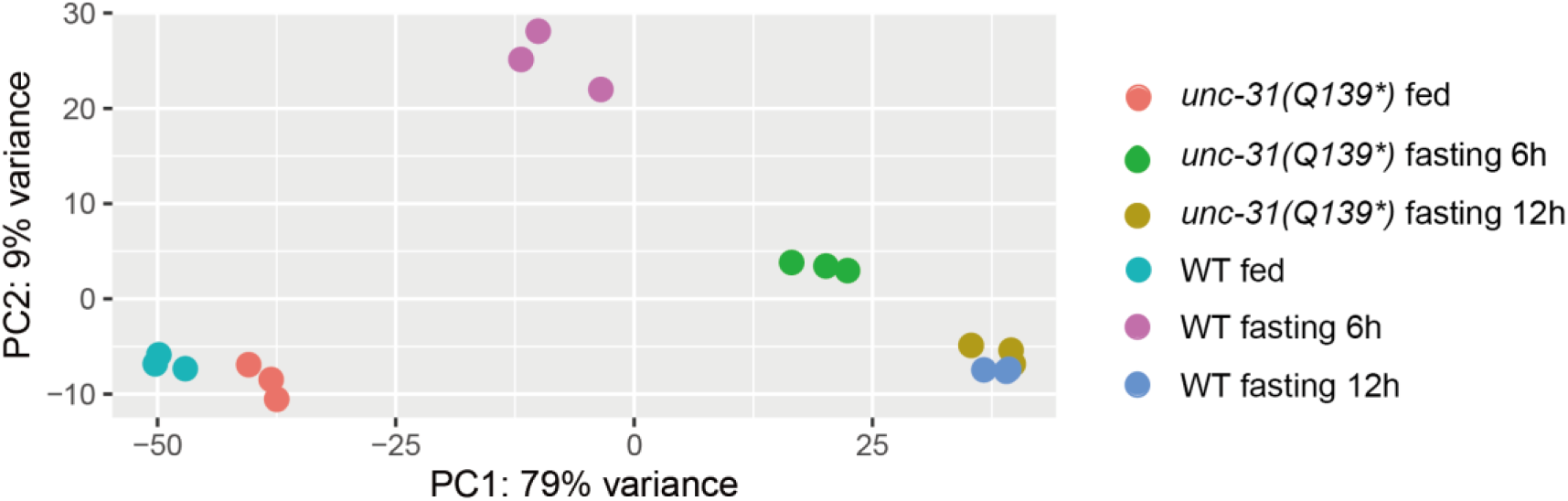
Principal component analysis (PCA) of RNA-seq from WT and *unc-31(Q139*)* mutants. N = 3.

**Figure S4.**
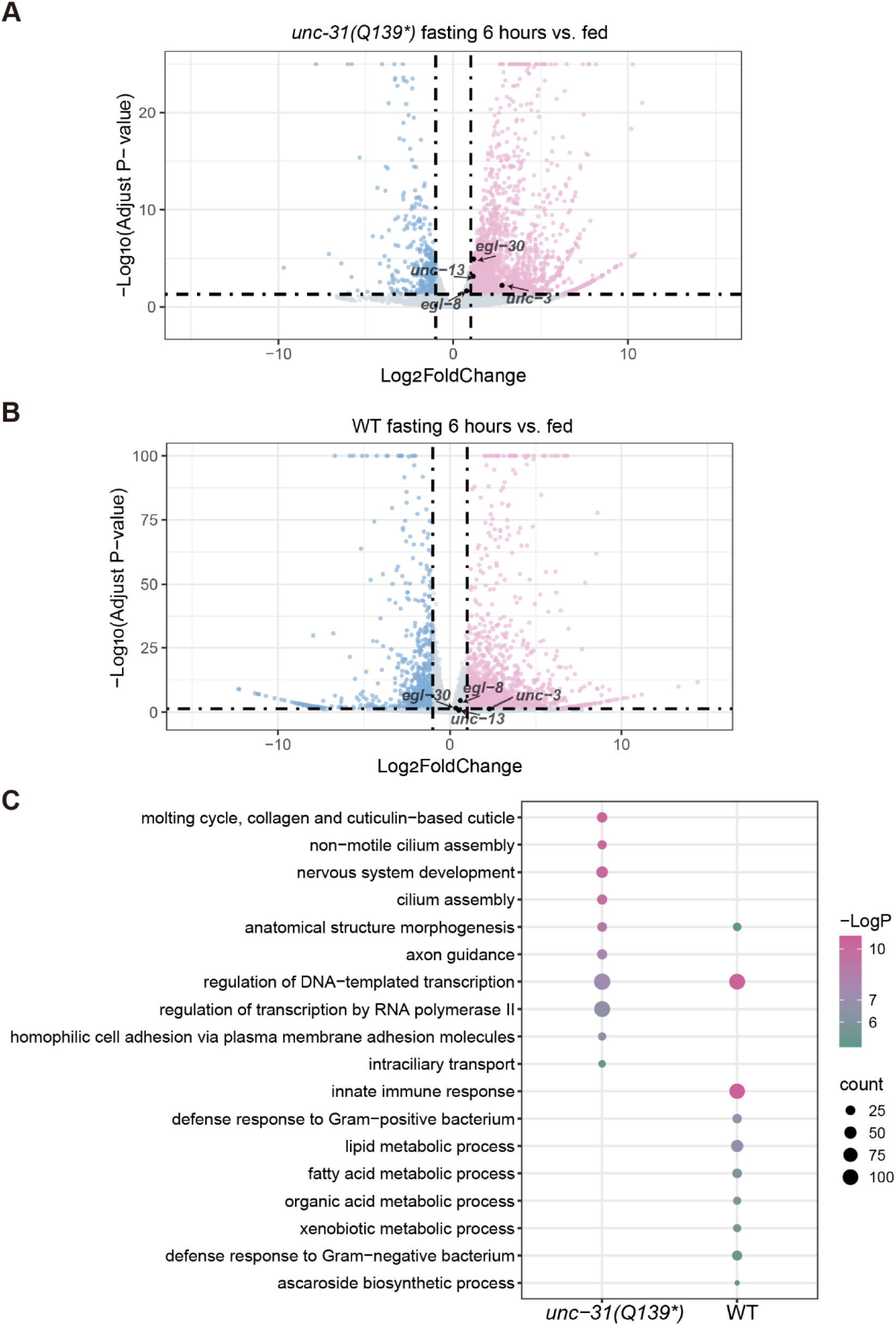
Transcriptomic profiling of *unc-31(Q139*)* mutants and WT animals following 6-hour fasting. **(A-B)**Volcano plots of deferentially expressed genes in *unc-31(Q139*)* mutants **(A)** and WT animals **(B)** after 6-hour fasting. Pink and blue dots represent genes that are significantly upregulated and downregulated, respectively (|fold change| > 2, adjusted P-value < 0.05). For clarity, points with -Log_10_(Adjusted P-value) > 25 (A) and > 100 (B) were capped at 25 and 100, respectively. **(C)** Dot plot of Gene Ontology (GO) enrichment analysis for biological processes in genes upregulated after 6-hour fasting in *unc-31(Q139*)* and WT animals. Dot colors indicate P-values and dot size corresponds to the number of genes enriched in each biological process.

**Figure S5.**
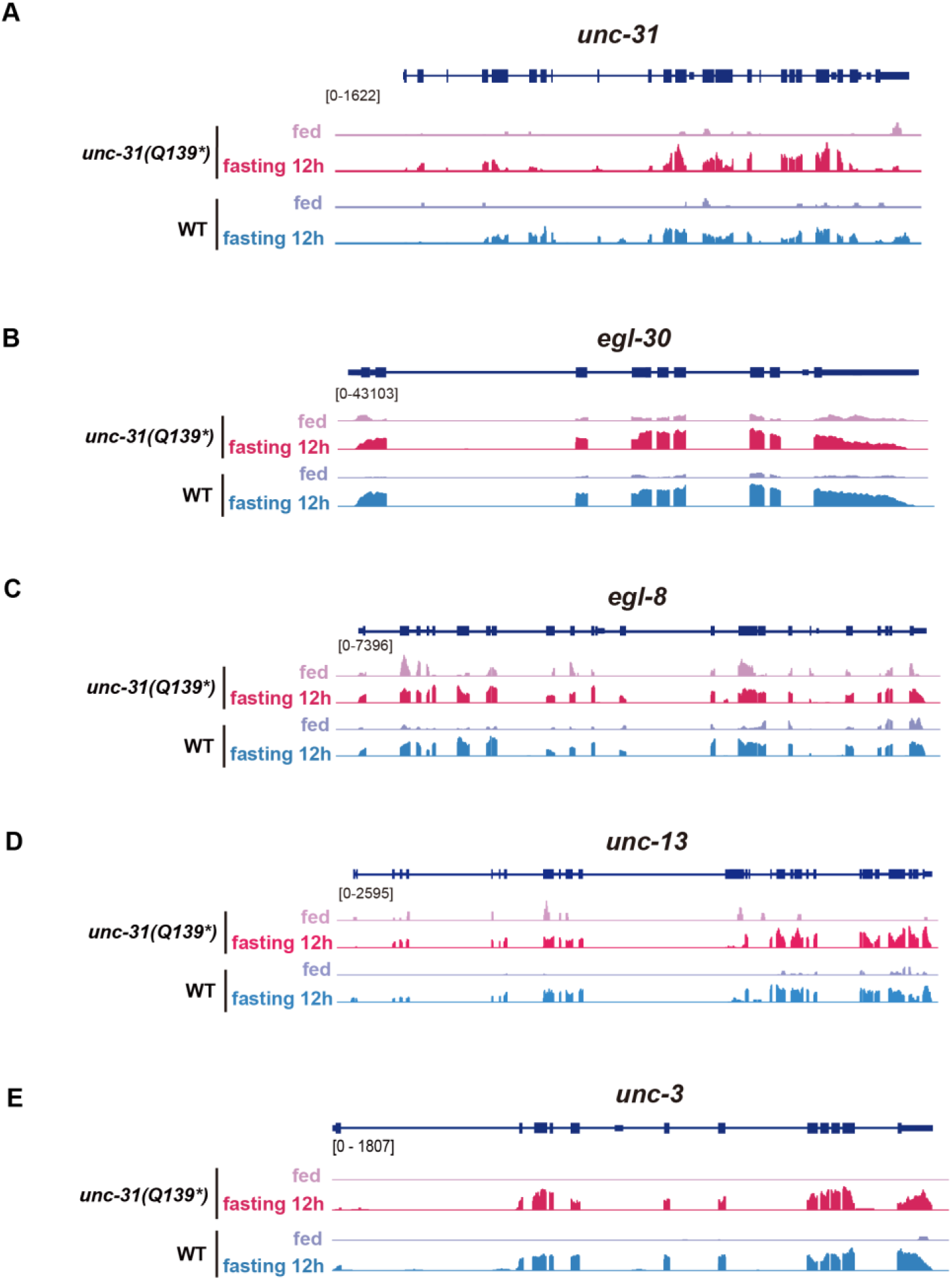
12-hour fasting induced upregulation of *unc-31*, *egl-30*, *egl-8*, *unc-13* and *unc-3.* **(A-E)** IGV visualization of distributions of normalized RNA-seq reads at *unc-31*, *egl-30*, *egl-8*, *unc-13* and *unc-3* loci in WT and *unc-31(Q139*)* mutants under fed and 12-hour fasting conditions.

**Figure S6.**
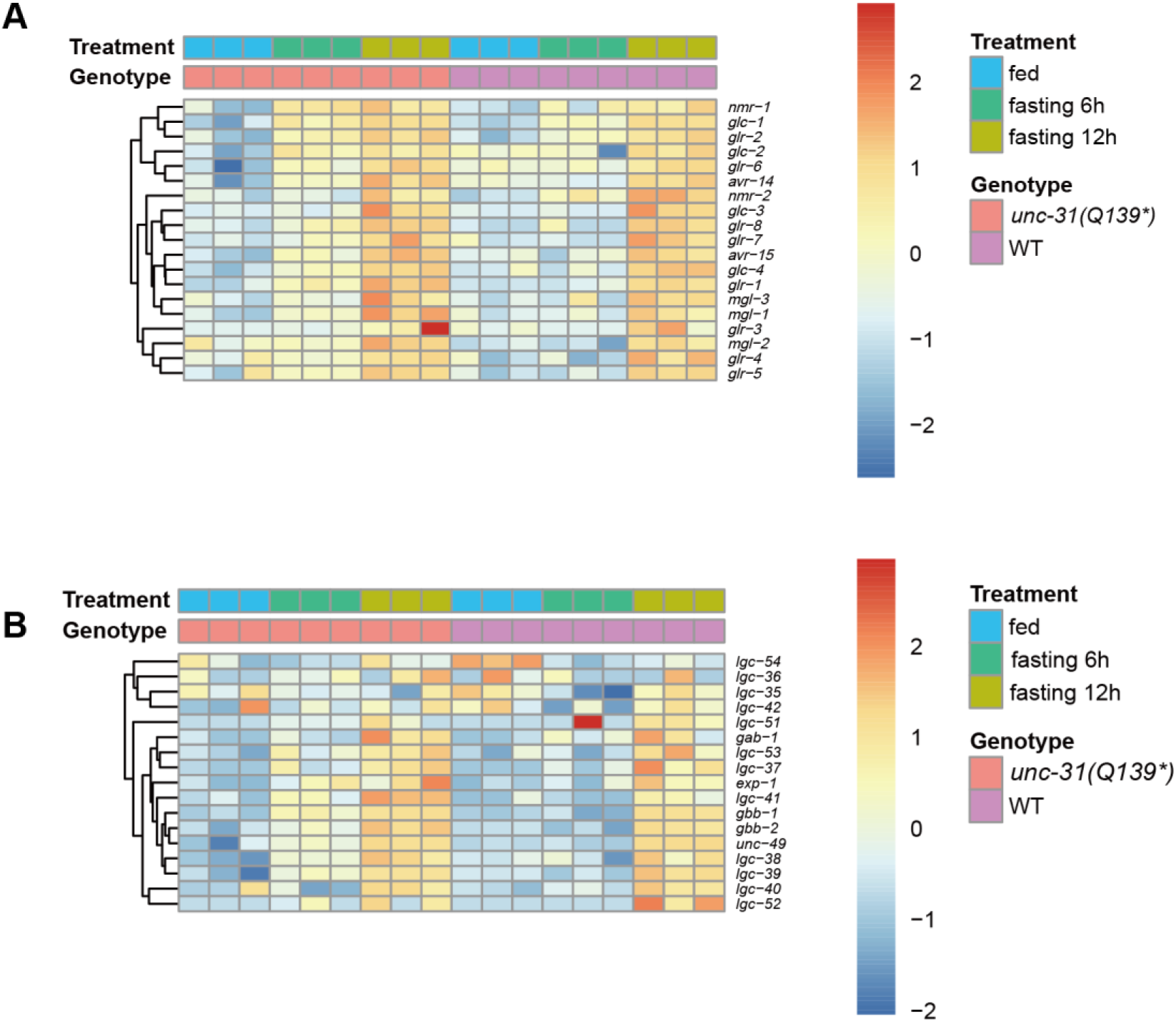
12-hour fasting induced upregulation of genes encoding glutamate receptors and GABA receptors. **(A)** Heatmap of expression levels of genes encoding glutamate receptors. **(B)** Heatmap of expression levels of genes encoding GABA receptors,

**Table S1.** Strains in this study.

| Strain Name | Genotype | Method |
| --- | --- | --- |
| *cas30000* | *unc-31(Q139\*)* | EMS screen |
| *CB169* | *unc-31(e169)* | (Brenner, 1974) |
| *cas30038* | *unc-31(Q139\*);egl-30(A227V)* | EMS screen |
| *cas30043* | *unc-31(Q139\*);Y34B4A.4(E445\*)* | EMS screen |
| *cas30044* | *unc-31(Q139\*);glt-1(Δ328-392)* | EMS screen |
| *cas30046* | *unc-31(Q139\*);egl-30(R25C)* | Genetic cross |
| *cas30047* | *unc-31(e169);egl-30(A227V)* | Genetic cross |

